# Exotic Hobos: Release, escape, and potential secondary dispersal of African red-headed agamas (*Agama picticauda* PETERS, 1877) through the Florida railway systems

**DOI:** 10.1101/2020.05.11.089649

**Authors:** Russell J. Gray

## Abstract

**Context:** Invasive reptile species are introduced and established through a variety of distribution channels (e.g. accidental/intentional release by pet owners, hitchhiking on imported goods), and can be detrimental to native ecosystems. Understanding the origins and spread of non-native species can help land managers to make informed decisions when attempting to prevent future introductions and remove established populations.

**Aims:** The objectives of this study were to employ modelling techniques with open-source observational data to confirm putative local origins of African red-headed agamas in Florida and to locate potential distribution channels in which they are spreading throughout the state.

**Methods:** Geographic profiling, a technique commonly used for criminal investigations, was used along with suspected origin locations of introduced African red-headed agamas (*Agama picticauda*) from the literature and observations from the Early Detection and Distribution Mapping System (EDDMapS). Anchor points and their immediate surroundings were investigated for potential patterns of origination and dispersal.

**Key results:** The results of this study provide evidence that African red-headed agamas likely established themselves through both intentional releases and unintentional escapes from the pet owners and breeders, while also potentially hitchhiking on plant exports, and dispersing throughout Florida via the railway systems.

**Conclusions:** Given the potential for railways as a method of assisted dispersal, and given the potential exports they may be hitchhiking on, the results of this study suggest that railcars and railway export facilities should be included in future management of non-native African red-headed agamas in Florida.

**Implications:** The implications of this study builds on prior evidence that geographic profiling is an effective modelling tool in regards to biological invasions, which can accurately confirm suspected origins while also effectively mapping previously unknown epicenters of population clusters to be investigated by managers.

## INTRODUCTION

Non-native species are often introduced to new areas unintentionally through human-activities, including the trafficking of wildlife (Romagosa 2015). This is especially true with herpetofauna, where local populations are established, and able to travel long distances via human assisted dispersal, creating additional sub-populations and increasing spread rate (Liu et al. 2014). Invasive herpetofauna are often detrimental to native wildlife: naïve to predators, outcompeting native species, and causing potential cascading effects on the local ecosystems in which they inhabit (Shine 2010; Stuart et al. 2014; Wilson 2017). The state of Florida hosts an astounding array of non-native species which have been introduced predominantly through human activity, many of which are herpetofauna (Bartlett and Bartlett 1999; Butterfield et al. 1997; EDDMapS, 2012; Engeman et al. 2011).

Among the introduced herpetofauna in Florida is the African red-headed agama, which has become increasingly wide-spread and established throughout several counties (Enge et al. 2004). Previously thought to be *Agama agama africana* Hallowell, 1844 (Harris, 1964) or *Agama agama*, Linnaeus 1758 (Wilson and Porras 1983), phylogenetic study by Nuñez et al. (2016) revealed that the introduced Florida agama populations are more likely to be *Agama picticauda* (PETERS, 1877). The molecular analysis which provided verification of the species (Nuñez et al. 2016), also provided several other potential geographic origins for the agamas, and therefore more potential natural history facets to take into account for their ability to thrive and disperse.

Although molecular techniques are often deployed to locate the geographic origin of introduced species on a global scale (Heinicke et al. 2011; Lees et al. 2011), inferences from molecular data is limited in its ability to be applied to the management of invasive species (Fitzpatrick et al. 2012). Lack of accurate and abundant data can impair and delay the development of informed management decisions (Tyre et al., 2003; Faulkner et al., 2015). Unfortunately, in regards to biological invasions, response time is a crucial component of successful eradication attempts, while delayed action or inaction can result in potentially devastating consequences (Grantham et al., 2009; Martin et al., 2012). Given this paradigm of necessary response time and potentially lacking data, researchers have turned to modelling techniques such as geographic profiling to rapidly detect the activity centers of biological invasions (Faulkner et al. 2017; Papini et al 2013; Stevenson et al. 2012).

Geographic profiling is a technique which is used most often in forensic criminology to model the spatial locations of crimes and identify the origin location or “anchor point” from which a criminal is operating (Rossmo 1999; Rossmo et al. 2014). Since its original use, geographic profiling has been modified and applied to infectious disease models (Verity et al. 2014), ecology (Faulkner et al. 2015; Faulkner et al. 2017), wildlife criminology (Faulkner et al. 2018; Romanach et al. 2019), and biological invasions (Faulkner et al. 2017; Papini et al 2013; Stevenson et al. 2012), where locations of committed crimes are instead substituted with sites of infection, suspected poaching locations, and animal observations records. The geographic profiling algorithm uses coordinate locations of points of interest (e.g. locations of a crime), and through a Bayesian framework, clusters the coordinates to locate a suspected epicenter (e.g. location where a criminal operates) while providing an accuracy measurement called a “hitscore” for suspect locations (O’Leary, 2009).

Using introduced African red-headed agama occurrence records retrieved from the Early Detection & Distribution Mapping System or EDDMapS (EDDMapS 2012) throughout Florida and implement the geographic profiling framework, I sought out to answer the questions: 1) What can geographic profiling tell us about the putative local origins of African red-headed agamas in Florida, 2) How effectively can the models verify or disqualify suspected origins of introduction locations from the literature, from the earliest observations, and from personal communications 3) what can geographic profiling say about potential dispersal mechanisms based on the surrounding environment of modelled anchor points.

## METHODS

### Study Species

African red-headed agamas are semi-arboreal, and thrive in urban environments within both their native and introduced ranges (Akani et al. 2013; Enge et al. 2004; Luiselli et al. 2011; Yeboah, 1982). Although their diet is primarily insectivorous, they are opportunistic omnivores that have been documented eating fruit and discarded anthropogenic food items (Akani et al. 2013; Anibaldi et al. 1998; Chapman and Chapman, 1964; Harris, 1964; Ofori et al. 2018; Yeboah, 1982). Agamas often take shelter in nearby objects, crevices, and structures upon approach by humans, to avoid unfavorable weather conditions, and for protection against predation attempts (Enge et al 2004; Cloudsley-Thompson 1981).

The first records of agamas in Florida appeared as far back as the mid-1970’s with voucher specimens (LSUMZ 36647, UF 43490) collected from a population thought to be eliminated by the demolition and subsequent building of a railway system (Enge et al. 2004; Wilson and Porras, 1983). The re-emergence of agama populations throughout Florida took place around the late 1980s to early 2000s where some individuals were expected to have been released after breeding facilities or exotic pet shops were destroyed by hurricane Andrew, while others may have been escaped pets, or released into the wild intentionally (Enge et al. 2004). Currently, there have been observations of A. picticauda in 17 Florida counties (Figure 1); some of which are assumed to have been isolated incidents, others which are established sub-populations. Although many origins of the African red-headed agamas in Florida are either assumed to be specific locations, or have been communicated by confession of the responsible individual (Enge et al. 2004), there have been no attempts so far to use modelling methods for determining the accuracy of these assumed origins.

**Figure 1.**
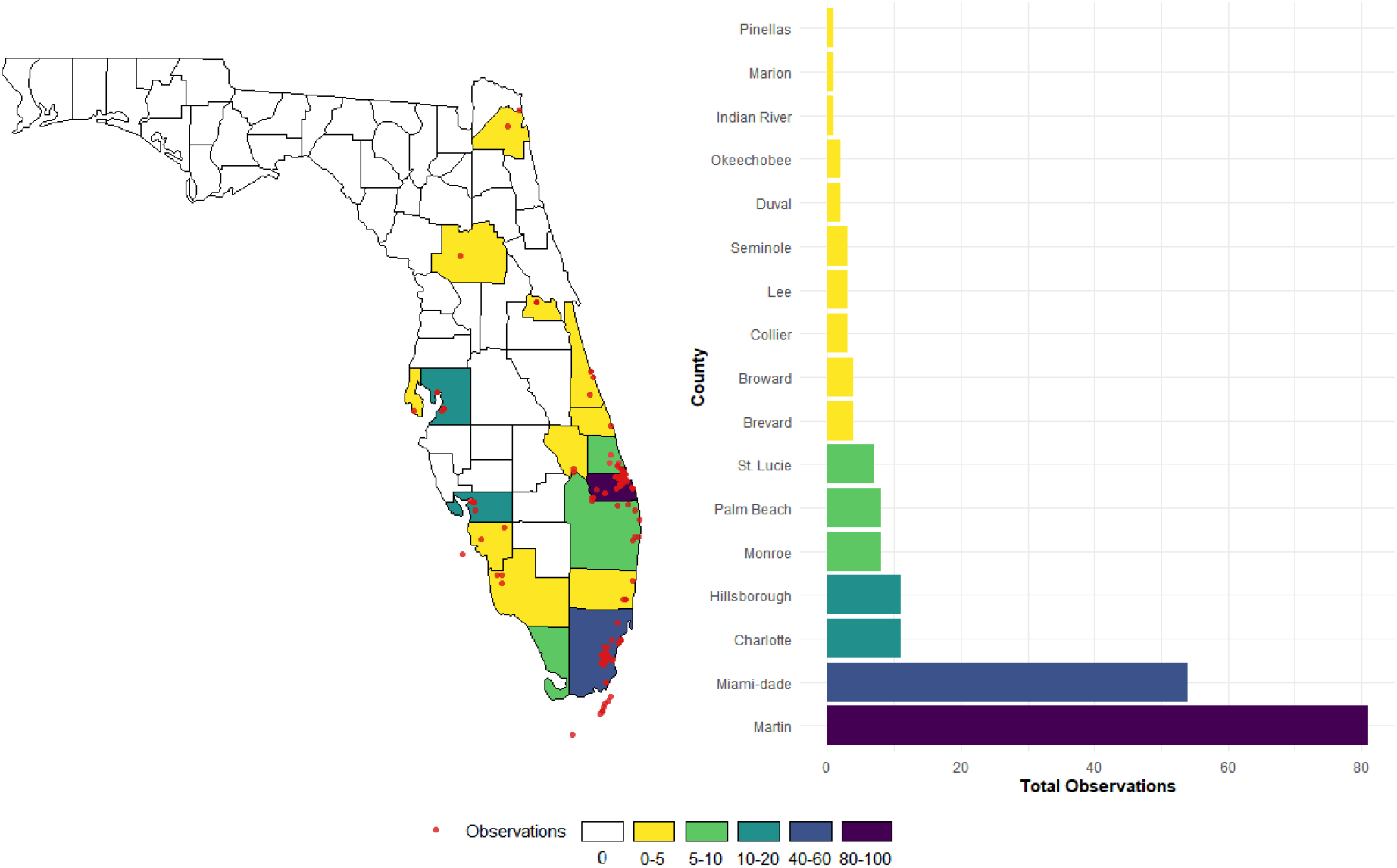
Total number of A. *picticauda* observations from EDDMaPS per Florida county between 2002 – 2015 (with the exception of a single 1975 specimen from Miami-Dade County; n = 194).

### Data

All records of *Agama picticauda* were queried from EDDMapS (EDDMapS 2012). Records with no coordinates were dropped from the data set, and coordinates which fell outside of the Florida state shapefile were masked from the data. After manual quality control checks of the data, 194 observations encompassed by 17 Florida counties remained and were used in the analysis (Figure 1).

### Analysis

All analyses were carried out using R statistical software version 3.6.3 (R Core Team 2020) and Google Earth Pro (Google n.d.). Suspected release locations (n = 6), as well as the earliest observation record (n = 1) (Table 1) were used in the geographic profiling model by employing the R package “RgeoProfile” (Verity 2015). The sigma mean (i.e. the mean prior standard deviation of dispersal distribution in km) was set to 0.5, as was the noted dispersal distance of a >10-year sustained population by Enge et al. (2004). All other parameters were set to the function default with weakly informative priors. Prior and posterior distributions were plotted and analysed for abnormalities before proceeding. HitScores (HS) were recorded for each suspect location; higher HS indicate low probability of the suspected origin, while lower HS indicate the suspected location is more likely to be the true origin site. Centroids of suspected origin areas were analysed on Google Earth Pro within a 5 km radius to examine if there were any evident trends in the surroundings (e.g. businesses, housing, land use, transportation routes, etc.). Euclidean distance from modelled locations to relevant points of interest were calculated in a post-hoc analysis to categorize modelled origin using the Google Earth Pro ruler tool. After the initial analysis, railway shapefiles were downloaded from the US Bureau of Transportation Statistics (BTS 2020). Other auxiliary R packages were used for data organization and visualizations (Supplementary file I)

**Table 1.** Locations of all suspected origins used in the geographic profile model. An asterisk * was placed next to locations used in the south Florida subset model.

| **Suspected origin** | **Longitude** | **Latitude** | **Year** | **Authority** |
| --- | --- | --- | --- | --- |
| **Reptile dealer residence \*** | -80.4556 | 25.53618 | 1992 | Enge 2004 |
| **Reptile dealer relocated** | -80.3199 | 27.13011 | 1999 | Enge 2004; powell, 2003 |
| **Coral castle \*** | -80.4443 | 25.50054 | 1992 | Pers. Comm. |
| ***A. picticauda* (Earliest observation)** | -80.3114 | 25.837 | 1975 | EDDMapS |
| **Strictly reptiles** | -80.2195 | 26.04573 | 1989 | Enge 2004 |
| **Private residence** | -82.0305 | 26.94154 | 1987 | Clark 2003 |
| **Released from reptile store** | -81.1684 | 28.4618 | 2000 | Enge 2004 |

Due to interesting patterns of anchor point locations (abundance of proximal exotic plant exporters), a subset of the total observation data was created to only include records from south Florida below decimal latitude 27.0° N. The subset of data was then divided into two groups: group one included observed locations (n = 49) between 2002 and 2015 (with the single voucher record from 1975), group two included the locally suspected origin locations: a) Coral Castle (Pers. Comm.), and b) the location of a reptile dealers’ home which was apparently destroyed in hurricane Andrew (Enge 2004). The model was run again with the subset and new modelled origin locations were analyzed.

## RESULTS

### South Florida Geographic Profile

The south Florida geographic profile identified Coral Castle (HS of 2.712) and the destroyed reptile breeders house near the junction of Coconut Palm Drive (SW 248th Street) and SW 163rd Avenue (HS of 0.190) as the primary suspected origin locations, which indicated the most likely origin of African redheaded agama sub-populations in south Florida to be the destroyed reptile breeders house. Due to the prominence of several sub-populations without connectivity, the model produced 18 potential origin locations (Figure 2). Upon examination of areas surrounding the location outputs, there was an evident trend in nurseries, importers, and exporters of exotic plants being within 0.5 km of the suspected origin locations (72%), although many were categorized as railway locations if they were proximal to railways. Two of the points were suspected escape/release locations, six points were identified as exclusive exotic plant locations, over half of the points fell within railway locations (~1 km or less from railroads; n = 8).

**Figure 2.**
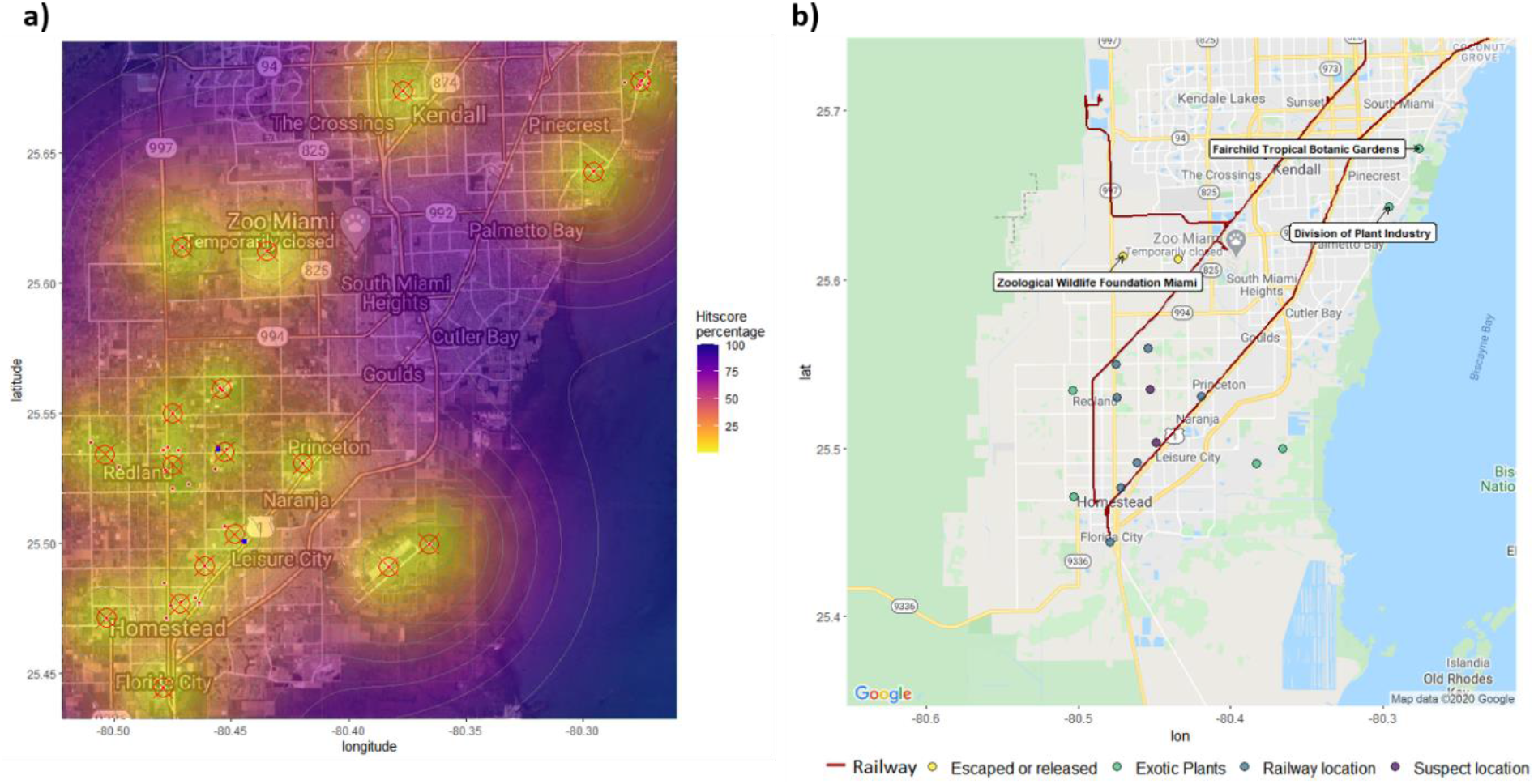
a) Geographic profile origin location output of African red-headed agamas (red crossed circles) in south Florida. Suspected origins (blue) include Coral Castle, and the location of a reptile dealer whose house was destroyed in hurricane Andrew; b) modelled origin locations categorized by relevant surroundings, suspect location matches, and labelled points of interest.

### Overall Geographic Profile

The overall geographic profiling model generated 46 origin locations for the sub-populations of A. *picticauda* throughout Florida, with an immediately evident trend of proximity to the Florida railway systems and an apparent outward dispersal from metropolitan areas being the most recent records (Figure 3). Out of the modelled locations, three locations were a complete match with the suspected origins: 1) the reptile dealer, whose house was suspected to have been destroyed during hurricane Andrew (HS = 0.002), 2) the earliest observation of A. *picticauda* at the Inter-Rail Transport of Miami HS = 0.086, and 3) the release at the private residence in Punta Gorda (HS = 0.003). All other suspected locations were closely matched with the modelled origin locations (Relocated reptile dealer, HS = 1.393; Coral Castle, 0.982; Strictly Reptiles, HS = 0.023), except for the suspected location of the Orange County reptile store release provided as a personal communication in Enge et al. (2004) (HS = 39.1558).

**Figure 3.1.**
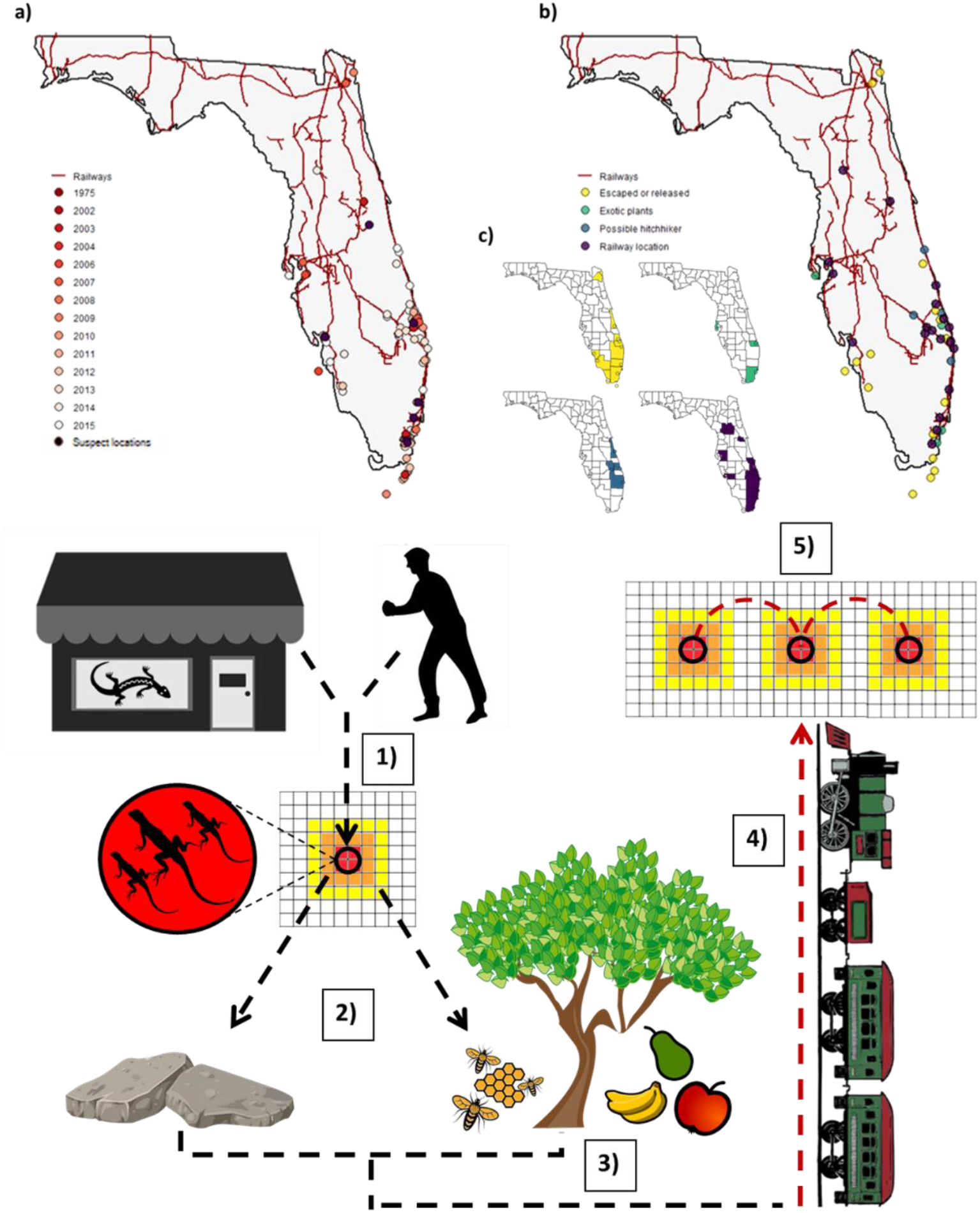
**(Top)** a) all observations of A. *picticauda* used in the model (n = 194) colored by associated year of observation (white = recent record; dark red = older record), with locations of the seven suspected origins (black); b) origins outputted by geographic profile (n = 46) model colored by category c) Florida maps with counties filled based on diaspora of each category. **Figure 3.2 (bottom)** Plausible origin and dispersal scenarios based on the data: 1) Agamas are released or escape from exotic reptiles shops/breeders or pet owners; 2) Agamas disperse to immediate debris pile shelter sites and then further disperse to allocate resources (e.g. as exotic plant exporters with pollinator prey); 3) rocks, cement products, and tropical fruits are loaded on trains and exported to warehouses around Florida; 4) Agamas hitchhike on the exported products to new locations.

Trends in modelled origin locations were divided into five categories: 1) Escaped or released (locations proximal to pet shops, reptile breeders/rescue centers, and locations which had no association with the other categories; n = 16); 2) Exotic plants (locations that were <1km from exotic plant nurseries, importers, or exporters; n = 4); 3) Possible hitchhiker (2-5 km from railroads, which assumedly travelled from the railroad to the current location via assisted dispersal; n = 4); 4) Railway locations (locations proximal to railways, most points were < 2 km from railways; n = 19); and 5) Suspect locations (locations which fell <1km of their suspected origin; n = 7). Railroad locations had a mean Euclidean distance of 581.96 m, with a maximum distance of 1.98 km, and a minimum of 13.27 m (Many agama observations themselves were directly over the railroad when these locations were produced). Mean Euclidean distance from suspect location matches was 388.38 m, with a max of 691.02 m (location proximal to the homestead reptile breeder’s house which was destroyed by hurricane Andrew), and a minimum of 0 m; the original observation of A. *picticauda* from the Inter-Rail Transport of Miami was not clustered by the model and was therefore outputted as a modelled location. All exotic plant locations were <0.5 km from the modelled origin location with agama observations occurring within the exotic plant importers/exporters or surrounding them.

## DISCUSSION AND CONCLUSION

By utilizing the geographic profiling method, a total of 64 locations were identified as potential origin sites for sub-populations of the invasive African red-headed agama in Florida (46 for the overall model; 18 for the south Florida model) The difference in the model output was expected, since the clustering algorithm varies based on spatial extent and occurrence points fed into the model. Each modelled location allowed me to evaluate potential trends in the surrounding environment and to verify the accuracy of suspected origin locations from the literature. This method precisely confirmed suspected origin locations and provided insight into how non-native species such as African red-headed agamas are likely dispersing throughout much of the state.

### Origins

African red-headed agamas were documented in south Florida as early as 1975, the first of which was found proximal to what is now the Inter-Rail Transport of Miami (Enge et al. 2004; Wilson and Porras 1983; EDDMapS, 2012). Given the typically small dispersal rate of the known sub-populations (~0.5 km per >10 years), the agama’s ranges aren’t likely to be greatly expansive from their points of origin via unassisted dispersal. For this reason, it is possible that the initial population in 1975 was not extirpated, but rather persisted unmanaged, allowing some individuals to be dispersed through the newly constructed railway to other locations. This could at least explain the widespread ranges in South Florida which aren’t proximal to the suspected release location in the Redlands. That said, there may not have been only a single introduction of Agamas into Florida, and there are other means for populations to establish themselves over long distances. After the phylogenetic verification by Nuñez et al. (2016) which built on the list of countries encompassing the Agama’s native ranges by Enge et al. (2004), it was concluded that the genetic variation in Florida sub-populations were indicative of multiple introductions. The results of this study also enforce the theory of multiple introductions, while contributing several potential assisted dispersal methods.

The geographic profiling models confirmed that the suspected origins of at least a few sub-populations in Florida were the reptile dealers’ home in Redlands, which was destroyed during hurricane Andrew, as well as the location of a reptile store near Hollywood, Florida. Additionally, there was a high model accuracy over the suspected private residence in Punta Gorda, which was reported to have released several individuals by Enge et al. (2004). However, there were sub-populations which fell somewhat far away from each of these suspected origins. One suspect location in south Florida fell within close proximity to the Zoological Wildlife Foundation (ZWF) Miami Venue, which has been operating since 2001 (Florida Department of State 2020). While this location is relatively distant from any railways when compared to the majority of other potential origin locations, it houses exotic reptile and plant species, which likely explains how the lizards originally arrived at this location and surrounding areas. Another modelled location fell between several observations directly on top of Central Florida Reptiles and Rescue in North Harbor City, although the observations themselves were lining the railways. Whether or not this facility participated in their spread is uncertain; still, it was interesting to see as the epicenter of the occurrences generated by the model. Furthermore, several points in various counties were scattered with single observations on houses and locations which were away from railroad tracks and exotic plant importers/exporters, which indicates these may have been released intentionally. It has been documented that some reptile owners will release lizards to build populations to capture and sell later on (Krysko et al. 2003). However, given that agamas tend to do poorly in captivity (Harris 1964), and are relatively inexpensive to purchase (Enge et al. 2004), some pet owners may simply release the animals to avoid caring for them.

Interestingly enough, while inspecting the immediate areas surrounding the south Florida model, one anchor point fell over the top of the sorting facility of the Florida Department Agriculture and Consumer Services (Division of Plant Industry), which sorts through imported exotic plants and attempts to control insects and pests which may negatively affect native Florida species (FDACS 2020). This brings into question whether there was an incident at the facility where African red-headed agamas hitchhiked on plants imported from countries where the species naturally occurs, and subsequently escaped, or whether it reached the facility by hitchhiking on a vehicle from a population already based in Florida.

### Dispersal Methods

Based on the anchor points produced by geographic profiling models, south Florida observations appear to be connected to areas with nearby exotic plant nurseries/exporters and railways outside of the suspected ranges of introduction, confirming the likelihood of plant affiliated secondary dispersal. Furthermore, nearly all of the recorded observations outside of major metropolitan areas occurred within close proximity to railways, or in shipping yards and warehouses in which they pass through. Also, worth noting is that observations in metropolitan areas typically occurred earlier (<2012) than the more recent observations (>2012) outside of them.

Due to their ambiguous diet (Akani et al. 2013; Anibaldi et al. 1998; Chapman and Chapman, 1964; Harris, 1964; Ofori et al. 2018; Yeboah, 1982), and tendency to take shelter in various structures (Enge et al. 2004; Cloudsley-Thompson 1981), it makes sense that the agamas would unintentionally hitch rides on the railways to other locations. Some notable areas in which the model pinged as the epicenter of observations along the railways included Cemex factories, tropical fruit exporters, exotic plant exporters, and grocery warehouses. The Cemex factories likely picked up hitchhiking individuals sunning themselves or sheltering in their product shipments. African red-headed agamas often tend to utilize man-made debris for perching and shelter (Enge et al. 2004; Grandison, 1968). Fruit exporters, grocery warehouses, and exotic plant exporters, on the other hand, have an abundance of insects (e.g. pollinators, flies, ants) which may subsist on their shipped and discarded product which creates a sustainable food source for the agamas. South Florida also has several bee farms which fall within close proximity to some of the agama sub-populations; one homestead bee farm near the railway has a thriving population (Pers. Obs.). Interestingly enough, the northernmost observations near Jacksonville also fall within close proximity to bee farms. Whether or not this is coincidence is uncertain; however, agamas being imported with bee farm related products is also a feasible explanation for the occurrences. Molecular sampling may reveal which sub-population they arose from.

### Effectiveness of Geographic Profiling

Geographic profiling has proven to be an effective method in generating potential epicenters of biological invasions (Faulkner et al. 2017; Papini et al 2013; Stevenson et al. 2012), with the advantage of performing well even with small amounts of data (Rossmo 1999) which can assist in rapidly responding to and mitigating non-native introductions early on. Compared to traditional kernel models, geographic profiling algorithms performs well with both single source as well as multiple source populations, which is often the case with biological invasions (Stevenson et al. 2012). The limitations of geographic profiling with occurrence data should be noted, as the clustering technique cannot identify clusters with relatively isolated points. Some of the suspected origin locations in the models were produced as the occurrence location itself in cases of coordinate isolation, which means that inferences were made based on the surroundings of the occurrence point itself and not necessarily an anchor point generated by a cluster of multiple points. Also, the use of observation data may not be suitable for species with large home ranges, as they may move further than their points of introduction in order to find more suitable conditions. That said, the models will work more accurately for species with small home ranges, and sessile species such as plants and fungi. In conclusion, geographic profiling is an invaluable technique which can be applied over a wide range of ecological investigations in order to locate potential points of origin and persons of interest (Faulkner et al. 2015; Faulkner et al. 2017; Faulkner et al 2018; Romanach et al. 2019; Verity et al. 2014). These models can be used by both ecologists and law enforcement agencies to generate a list of potential suspects to investigate, and create risk-management guidelines to effectively combat the spread of established invasive species and predict potential high-risk areas for the prevention of future introductions. More importantly, species observation records have been shown to uphold accuracy in geographic profiling models in comparison to the expensive, and otherwise time-consuming trapping approaches (Faulkner et al. 2017), which outlines the importance of open citizen-science data, and the employment of effective modelling techniques with minimal data.

### Implications

The implications of the findings in this study are not only applicable to Florida, as African red-headed agamas appear to be occupying a mode of transportation which is connected nationwide, and not only applicable to the species in question, as others species have the potential to hitchhike on shipped items and travel in the same manner (Rochford et al. 2015). Furthermore, railways and other transportation corridors are known sources of travel and establishment of invasive species populations (Van Der Windt and Swart 2008). Along with globalization of trade and shipping, invasive species being imported as stowaways (Hulme 2009) and through the pet trade (Perrings et al. 2005) becomes increasingly more prominent. Although the survival and spread of African red-headed agamas in other states may rely on their thermal tolerance, the structures and debris they use for shelter may allow them to survive cold temperatures for prolonged periods of time (Enge et al. 2004).

Considering the fruit exports, such as citrus, from Florida to other states and even other countries, the potential of agamas establishing themselves at warehouse locations along railways and ports of entry is increasingly likely. In order to effectively manage the spread of African red-headed agamas and other hitchhiking invasive taxa within their potentially habitable ranges, sub-populations within proximity to railways and ports of entry with transportation corridors should be targeted and effectively removed.

## Acknowledgements

I would like to extend my gratitude to those who operate and contribute to EDDMapS and all other open science outlets which allow researchers to freely analyze and discover trends in data. I would also like to thank Cameron Hodges and Eva Gazagne for their helpful review, suggestions, and edits to the manuscript.

